# Benchmarking computational tools for de novo motif discovery

**DOI:** 10.1101/2024.01.12.574168

**Authors:** Leandro Simonetti, Ylva Ivarsson, Norman E. Davey

## Abstract

**Background:** Over the past twenty years, numerous motif discovery bioinformatic tools have been developed for discovering short linear motifs (SLiMs) from high-throughput experimental data on domain-peptide interactions. However, these tools are generally evaluated individually and mostly using synthetic data that do not accurately capture the motif context observed within proteomic data. Consequently, it is unclear how these tools perform in real-world use cases and how they perform compared to each other.

**Results:** Here, we benchmarked five motif discovery tools and seven general sequence alignment tools on their capacity to find SLiMs. For this purpose we have built MEP-Bench, a benchmarking dataset of peptides of varying complexity from curated SLiM instances from the Eukaryotic Linear Motif database. MEP-Bench allows tools to be tested for the effect of dataset size, peptide length, background noise level and motif complexity on motif discovery. The main metric used to compare all tools was the percentage of correctly aligned SLiM containing peptides. Two motif discovery tools (DEME and SLiMFinder) and a sequence alignment tool (Opal) outperformed the rest of the tools when benchmarked with this metric, averaging over 70% correctly aligned motif-containing peptides. The performance of the motif discovery tools and Opal were not affected by the sizes of the datasets. However, increasing peptide lengths and noise levels decreased all tools’ performances. While all tools performed well for N-/C-terminal motifs, for low-complexity motifs only DEME and SLiMFinder returned correctly aligned motifs for 50% or more of the datasets.

**Conclusions:** This study highlights DEME, SLiMFinder and Opal as the best performing tools for finding motifs in short peptides, and it indicates experimental parameters that should be considered given the limitations of the available tools. However, there is room for improvement, as no tool was able to identify all motif types. We propose that MEP-Bench can serve as a valuable resource for the SLiM community to compare new motif discovery methods with those benchmarked here.

## Background

Protein-protein interactions underlie cell function and elucidating them is key to understanding how cellular systems work. Many of these interactions are mediated by compact short linear motifs (SLiMs), which are typically 3-10 amino acid stretches found in the intrinsically disordered regions of proteins (1). SLiMs play major roles in cell signalling, protein trafficking and homeostasis and it has been estimated that the human proteome contains about 100,000 SLiMs (1). Less than 5,000 instances are currently annotated in the Eukaryotic Linear Motif (ELM) database (2), which is a gold standard database for SLiMs.

Significant experimental efforts are being invested in finding SLiM-containing peptides. The experimental methods include peptide arrays coupled to mass-spectrometry, phage display of peptides from given proteomes, as well as functional assays such as transactivation assays and degradation assays (3). These methods typically generate a set of peptides that directly or indirectly bind to a given protein or exert a given function. Depending on the experimental design, the complexity of the results and the number of output peptides a consensus binding motif can often be generated. This is a key quality control step for these discovery methods as an enriched motif generally suggests that specific and biologically relevant binding events have been observed.

For some analyses, motif discovery can be a trivial challenge, especially if the experiment returns clean data with little background noise, and the interrogated bait recognises a highly defined type of SLiM. For example, many WW domains specifically bind to PPxY containing peptides (**Figure 1A**), a SLiM that has been established using many different techniques including peptide SPOT array (4), yeast two-hybrid (5), and peptide-phage display (6). Other types of SLiM-binding proteins produce more complex data, such as calmodulin, an EF-hand protein that recognises aliphatic helices with variable spacing between the key hydrophobic residues that dock into the binding sites (7,8) (**Figure 1B)**. Finally, post-translational modification sites tend to have very short and low complexity motifs, like the S/TxxE motif recognised by casein kinase 2, a serine/threonine kinase that phosphorylates different targets and has an acidic residue 3 positions after the modification serine or threonine as its main specificity determinant (9–12) (**Figure 1C**). Analysing peptides containing such low complexity motifs is difficult regardless of the quality of the generated data.

**Figure 1.**
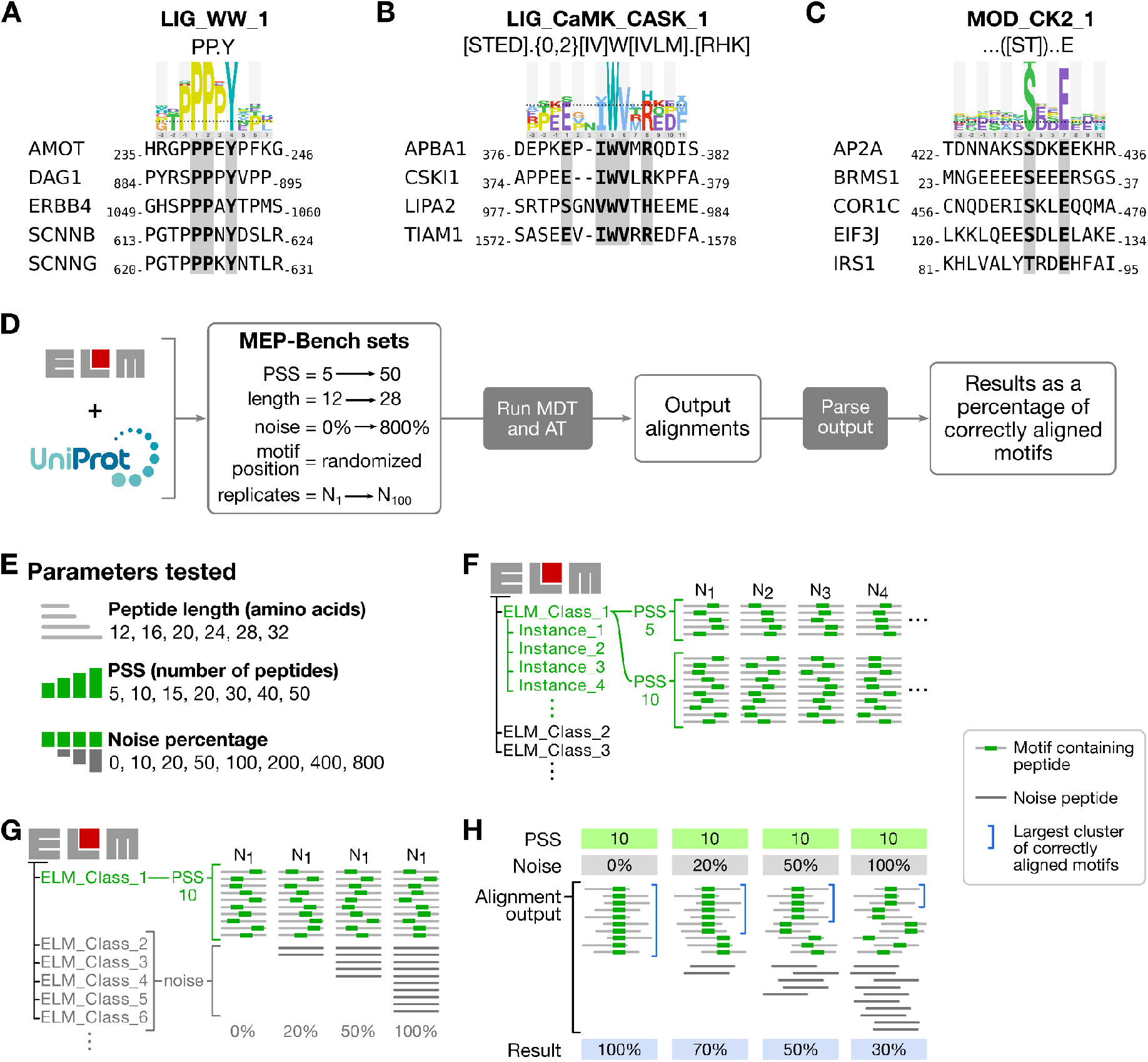
SLiMs and benchmarking. **(A-C)** ELM class names, regular expression definitions and logos are shown for the defined motifs recognised by WW **(A)**, calmodulin **(B)** and casein kinase 2 **(C)** domains, together with a selection of human peptidic sequences obtained from ELM instances for the respective classes to showcase the sequence diversity. **(D)** Overview of the benchmarking process. **(E)** Values tested for the three parameters evaluated in the current work. **(F)** Benchmarking datasets were generated from available ELM classes. From each ELM class, a number of instances (belonging to different unique proteins) were randomly picked according to the positive set size (PSS) parameter and, using the instance motif information, a peptide of the desired length including the motif in a random position was generated from the complete protein sequence in UniProt. This process was repeated to generate the desired number of replicates. **(G)** Different noise percentages were added to each benchmarking dataset by randomly sampling other ELM classes, generating a single peptide from a single random instance for each sampled ELM class. Noise percentages are defined on the base of the PSS, so 50% noise for a PSS of 10 would mean 5 noise peptides are added to the dataset. **(H)** The resulting output alignments from all motif discovery tool and alignment tool runs were scored as the percentage of correctly aligned motifs in the PSS peptides group. When multiple groups of correctly aligned peptides were found, the score corresponded to the biggest group. Logos in A-C are representations of PSSMs calculated using a relative binomial distribution for all instances present in each ELM class and the amino acid enrichments were calculated with the background being all human proteome disordered regions, the dotted line represents p=0.05, built using LogoViewer (21).

Numerous bioinformatic tools have been created over the years. For example MEME is an algorithm that searches for common significant non-gapped patterns in a set of input sequences and then scores them depending on their similarity and presence in the input dataset (13,14). SLiMFinder on the other hand can discover gapped motifs by combining amino acid dimers based on their abundances in the input data, building motifs from them and assigning them a significance score based on their probability of randomly appearing in the input dataset (15). Some methods were developed with specific experimental data in mind, like GibbsCluster, an alignment and clustering algorithm based on Gibbs sampling (16), developed with the motivation of identifying consensuses in sets of major histocompatibility complex binding peptides obtained via high-throughput microarray experiments. MUSI was also developed with results from high-throughput analysis in mind, using a mixture model to rapidly find multiple motifs in large datasets generated from microarray or phage-display followed by next-generation sequencing (17). DEME on the other hand was specially designed to reduce overfitting motifs in small protein datasets (18) and it does so by comparing a positive and negative set of sequences using a probabilistic model. Similarly, MotifHound and DALEL are methods that also rely on a negative or background set, as they are based on exhaustive enumeration of possible motifs in a dataset followed by an overrepresentation scoring of these motifs in a background proteome (19,20).

In this study, five widely used motif discovery tools and seven classic multiple sequence alignment algorithms were benchmarked (**Table 1**). For this objective, several benchmarking datasets, structured to emulate the output of high-throughput methods, were generated based on SLiM instances curated in the ELM database (**Figure 1D**). We call this collection of datasets the Motif Extraction from Peptides Benchmarking (MEP-Bench) sets. We evaluate different challenges encountered when analysing peptide data related to the number of sequences available, the length of sequences identified, the amount of noise and the complexity of the underlying motif. We discuss advantages and disadvantages of the different methods and provide guidelines for which method to use for different kinds of data and research questions.

**Table 1.** Motif discovery tools and alignment tools.

| Group | Tool name | Runs locally (OS)* | Web-tool | Used in study | Output | Notes | Refs |
| --- | --- | --- | --- | --- | --- | --- | --- |
| MDT | DALEL | Yes (W) | No | No** | Motif | Requires a background proteome | (20) |
| MDT | DEME | Yes (M/GL) | No | Yes | Alignment | Requires a negative set (background) and a motif length value | (18) |
| MDT | GibbsCluster | Yes (M/GL) | Yes | Yes | Motif + alignment |  | (16) |
| MDT | MEME | Yes (M/GL) | Yes | Yes | Motif + alignment |  | (13) |
| MDT | MotifHound | Yes (W/M/GL) | No | No*** | Motif | Requires a background proteome | (19) |
| MDT | MUSI | Yes (M/GL) | No | Yes | Motif + alignment |  | (17) |
| MDT | SLiMFinder | Yes (W/M/GL) | Yes | Yes | Motif |  | (15) |
| AT | Clustal Omega | Yes (W/M/GL) | Yes | Yes | Alignment |  | (22) |
| AT | ClustalW | Yes (W/M/GL) | Yes | Yes | Alignment |  | (23) |
| AT | MAFFT | Yes (W/M/GL) | Yes | Yes | Alignment |  | (24) |
| AT | MUSCLE | Yes (W/M/GL) | Yes | Yes | Alignment |  | (25) |
| AT | Opal | Yes (W/M/GL) | No | Yes | Alignment |  | (26) |
| AT | ProbCons | Yes (W/M/GL) | Yes | Yes | Alignment |  | (27) |
| AT | T-Coffee | Yes (W/M/GL) | Yes | Yes | Alignment |  | (28) |

List of all motif discovery tools (group MDT) and alignment tools (group AT) that were considered for the benchmark. The operating system for local execution, existence of a web-based portal, output result type (classified as alignment and/or motif) and original publication are shown for each tool as well as if they were used in the study. Extra notes are provided for those programs that require specific input. \*Operative systems are W: Windows, M: MacOS, GL: GNU-Linux. \*\*DALEL was excluded from this study because it does not run on GNU-Linux. \*\*\*MotifHound was tested but ultimately excluded for the study (see results section).

## Methods

An overview of the procedures performed to benchmark the different motif discovery tools (MDTs) and alignment tools (ATs) is shown in **Figure 1D**. The MEP-Bench sets were generated utilising curated motif instances from the ELM database (2) (**Figure 1E-G**), and five MDTs and seven ATs (**Table 1**) were run on each dataset. Notably, each tool provides different outputs, with most of them producing alignments of the input peptides together with one or more motifs in the form of a core consensus, as a position-specific scoring matrix (PSSM), or regular expression definition of the motifs. In order to allow the direct comparison, the data from all methods was converted to peptide alignments. The resulting alignments were then analysed and compared on the basis of the percentage of correctly aligned motifs (**Figure 1H**).

### MEP-Bench datasets generation

Several parameters were tested in the motif discovery benchmarking: (i) the length of the peptides (12, 16, 20, 24, 28, 32 amino acids); (ii) the number of peptides that contain the expected motif (i.e. the positive set size or PSS; 5, 10, 15, 20, 30, 40, 50 peptides); and (iii) the proportion of the peptides as a percentage of the PSS of the dataset that are noise (i.e. peptides that lack the expected motif, 0% (no noise) 10%, ∼20%, ∼50%, 100%, 200%, 400%, 800%) (**Figure 1E**). The true positive sets of peptides (number according to the PSS) were randomly picked from the list of possible instances belonging to 158 ELM classes (as of February 2020, **Supplementary Table S1**) (**Figure 1F**). We built datasets from each ELM class that contained the number of instances matching the PSS value. The datasets were built by selecting peptide sequences of the required length overlapping the ELM instance. The ELM instance was randomly positioned within the peptide and the whole motif was present in the final peptide. The positioning of peptides tiling N- and C-terminal motifs could not be randomised, as the free termini are required for these motifs. We ensured that each peptide only contains a single motif instance by excluding peptides with more than one non-overlapping hit for the ELM class regular expression. As each ELM class contains different numbers of curated instances this limited the number of datasets that could be generated for each PSS value (**Supplementary Figure S1A**), with the PSS of 5 spanning over 150 ELM classes while the PSS of 20 dropped to ∼35 and the PSS of 50 was only possible for a single ELM class (DOC_WW_Pin1_4). The number of usable ELM classes also varied slightly depending on the peptide length used (**Supplementary Figure S1B**), due on one side to some ELM classes containing instances with motifs longer than the peptide length value, and also as a result of enforcing a single expected motif per peptide, which excluded some instances from longer peptides. To limit the number of cases studied, the complete range of PSS values was only assessed for 16 amino acids long peptides, while for the rest of the peptide lengths only sets of 15 and 20 PSS were created.

To generate noise peptides in the datasets, peptides from other ELM classes were randomly added to the dataset (**Figure 1G**). Noise levels were defined as a percentage value of the PSS used for each dataset, so for example a 50% noise value for a PSS of 10 equals 5, while the same noise value for a PSS of 50 is 25. The absence of the PSS expected motif in the noise set was enforced by performing a search with the ELM class regular expression on the noise peptides. To account for the possible effects of all the randomised parameters including instances used, motif position in the peptides and noise peptides used, we generated 100 independent datasets (replicates) for each combination of PSS, length and noise values for each ELM class. Some noise values were skipped or slightly altered for specific PSS values and the highest noise level for PSS of 40 and 50 was limited by the total number of ELM classes available to 400% (**Supplementary Figure S1B**).

In total, approximately 650,000 benchmarking sets of peptides were created and the collection of all of these was named the “Motif Extraction from Peptides Benchmarking” (MEP-Bench) sets (**Supplementary Data SD1)**. The algorithm to generate datasets was written in Python.

### Motif discovery tools and alignment tools running and parameters

A Python wrapper was built to sequentially run all MDT and AT. The script was parallelised and run on all datasets utilising Python’s multiprocessing library. The benchmarking was performed in a Docker container running Ubuntu on a server with a 2 × 12 core Intel(R) Xeon(R) Silver 4116 CPU @ 2.10GHz with 64 Gb of shared RAM. All MDTs and ATs were run with their default parameters, except for the following detailed cases. For Clustal Omega an iteration value of 100 was used. For MEME a single motif was searched using the classic mode (no negative control provided). As DEME requires a motif length value and a negative control background set of peptides as inputs, the software was run multiple times with motif length values ranging from 3 to 13, and a 100 pure noise peptides (PSS = 0) dataset was generated for each assayed combination of ELM class and peptide length to use as the required negative background set. For MotifHound the background used was the curated disordered regions from the human proteome available from MobiDB (29). For GibbsCluster no insertion or deletion were allowed. For SLiMFinder, as it outputs a regular expression for a predicted motif and an alignment was required for scoring programs performances, the PSSM based alignment tool psiblastPSSM from the PSSMSearch package (30) was used to turn the output from SLiMFinder into the required output. Only tools that could run locally in a GNU-Linux system were used for the current work, which removed DALEL from the list of benchmarked MDTs (**Table 1**).

### Analysis and plotting

The percentage of correctly aligned motifs was calculated for each MDT and AT sequence alignment result as follows: All possible pairs of motif-containing peptides in a sequence alignment output were compared and a matrix was built where a value of 1 was assigned if the first position of the expected motifs in the peptide pair were aligned (that is, had the same index position including gaps), otherwise a value of 0 was assigned. The largest cluster of peptides with their motifs aligned was determined using scipy’s hierarchical clustering’s linkage function, and the number of peptides in this cluster was divided by the maximum number of motif-containing peptides (that is, the PSS of the dataset) to calculate the percentage of correctly aligned motifs (**Figure 1H**). As DEME produced 11 distinct output alignments (obtained from setting the motif window value from 3 to 13 amino acids), the clustering was performed separately on each of the 11 generated outputs, and the alignment with the highest percentage of correctly aligned motifs was selected for the further analysis.

For the ELM class level analysis, the motifs were classified as internal, N-terminal or C-terminal based on the motif consensus regular expression (**Supplementary Table S1**). ELM classes with motifs that could be both N- or C-terminal and internal were classified as terminal. ELM class motif probabilities were taken from the ELM database and binned into 3 “complexity” tiers: High (motif probabilities of <10^−5^), medium (10^−5^ to 10^−3^) and low (10^−3^ to 1).

For plotting purposes data was collapsed per tool by averaging their resulting percentage of correctly aligned motifs for all replicates and ELM classes datasets contributing to the combined set of PSS, noise percentage and peptide length parameters. All plots were built with the Seaborn and Matplotlib libraries for the Python v3.11 programming language.

## Results

Experimental methods aimed at exploring SLiM-based interactions typically produce a list of binding peptides as their output. Depending on the experimental set up, the datasets may contain direct biophysical ligands that contain a shared consensus motif, and also background noise caused by background binders (e.g. binding to plastic), by peptides giving false positive signals (e.g. due to autoactivation in yeast two-hybrid) or by otherwise non-specifically retained peptides. Finding the motif-containing peptides within a complex dataset can thus be a challenging task. Different computational tools have been generated for the purpose, and we here evaluated the performance of five of them together with seven general alignment tools (**Table 1**). To address which of these tools would be more suitable for analysis of experimental data, we generated the Motif Extraction from Peptides Benchmarking (MEP-Bench) sets, a series of datasets of varying complexity that emulate the output of high-throughput experimental methods.

### Selection of the motif discovery tools and alignment tools

From the seven motif discovery tools (MDTs) considered for the current work (**Table 1**), DALEL was excluded as it does not run natively on GNU-Linux systems. The remaining six MDTs and seven alignment tools (ATs) were then tested on a smaller subset of the generated datasets to test the pipeline. A test-run with 50 datasets replicas of the ELM class “LIG_Rb_LxCxE_1” was performed and the run times of each tool was evaluated (**Figure 2A**). The MDT “MotifHound” showed an average running time of 1,440 seconds (**Figure 2B**), 15 times slower than the next slowest tool (SLiMFinder) and for this reason it was removed, reducing the total number of MDTs evaluated to five. All alignment tools in **Table 1** were used from this benchmark.

**Figure 2.**
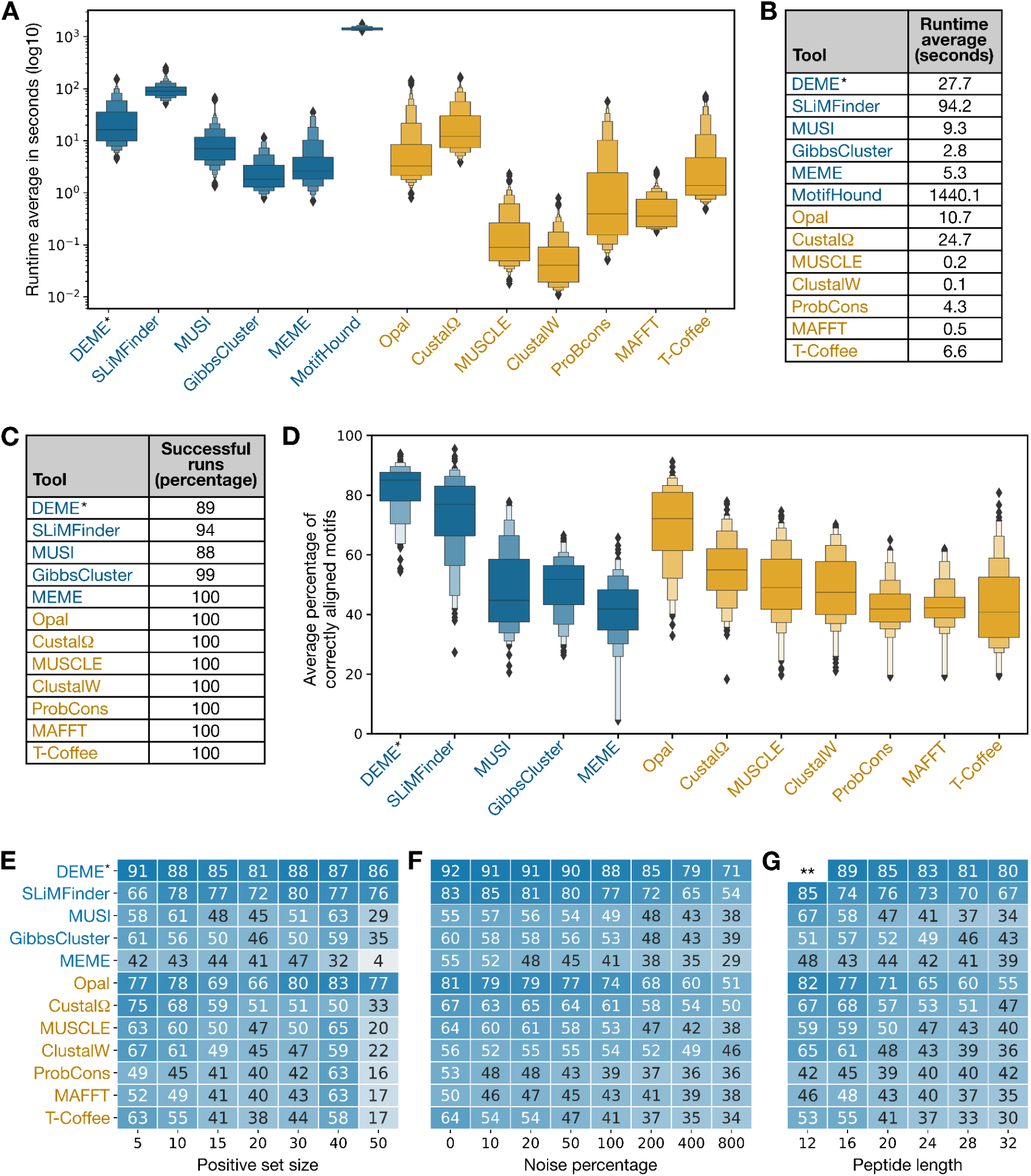
MDTs and ATs benchmarking results. **(A-B)** Running times in seconds for MDTs and ATs evaluated with a small fraction of the datasets. **(C)** Percentage of successful runs for MDTs and ATs on all datasets. **(D)** Overall results of the performance of all benchmarked motif discovery tools (blue) and alignment tools (orange). Each boxenplot shows all results by tool collapsed at PSS, peptide lengths and noise percentage level. **(E-G)** Effect of positive set size **(E)**, noise percentage **(F)** and peptide length **(G)** on the percentage of correctly aligned motifs by tool. The heat maps represent the average of all results for the respective tool and parameter values. The noise percentage groups for 20% and 50% also contain the tweaked values shown in **Supplementary Figure 1B**. *DEME results are the best results out of 11 runs using different expected motif length input values. **DEME failed to run on 12 amino acid long peptides producing results only for 2 replicates (**Supplementary Figure 2C**).

### Overall performance of motif discovery tools and alignment tools

The five selected motif discovery tools and the seven alignment tools were run on the complete MEP-Bench sets (**Supplementary Table S2**). The ATs produced an output for all datasets but for the MDTs, MUSI, DEME and SLiMFinder failed to return a result in more than 5% of the runs (**Figure 2C**). MUSI had trouble running on small datasets as only 47% of the runs for a positive set size (PSS) of 5 and 87% for a PSS of 10 produced results (**Supplementary Figure S2**). SLiMFinder shows a lower (73%) percentage of successful runs for a PSS of 40 and the reason behind this is harder to pinpoint though it seems to be related to the ELM classes in the set. Finally, DEME failed to run on 12 amino acid long peptides.

The motif discovery performance of the tools was evaluated by calculating the average percentage of correctly aligned motifs (**Figure 1H**). Initially, we assessed the overall performance of all tools (**Figure 2D**) and discovered that among the motif discovery tools, DEME and SLiMFinder outperformed the rest. It is crucial to note that DEME necessitates an input value for the motif size (alongside a negative background set), so it was executed 11 times and the optimal outcome was selected from among them. Surprisingly, we found that the alignment tool Opal ranked third in performance, surpassing other dedicated motif discovery tools.

We then investigated the impact of the parameters we evaluated (PSS, noise percentage, and peptide length) on the performance of the various tools (**Figure 2E-G**). Across the range of tested positive set sizes from 5 to 30 (as PSS=40 and 50 contain only 4 and 1 ELM class, respectively), all MDTs and the alignment tool Opal exhibited minimal to no impact of the PSS value on their average performance (**Figure 2E, Supplementary Figure S3**). This was also true for PSS values of 40 and 50 for DEME, SLiMFinder, and Opal. However, the remaining ATs did show a decrease in performance as the PSS increased. In contrast, the impact of added noise (**Figure 4C, Supplementary Figure S4**) and peptide length (**Figure 4D, Supplementary Figure S5**) was consistent across all tools: As they increased, all tools reduced the percentage of correctly aligned motifs.

Overall, three tools emerged as the top performers: The motif discovery tools DEME and SLiMFinder and the alignment tool Opal. Increasing the size of the datasets, either by adding more motif-containing peptides (larger PSS), introducing noise, or employing longer peptides, had the expected impact on alignment tools (except for Opal). Specifically, the percentage of correctly aligned motifs decreased as the search space expanded. This trend was also observed for MDTs with increased noise and peptide length, but not PSS. MDTs exhibited a relatively stable performance regardless of the number of motif-containing peptides.

### Impact of motif position and complexity

As motifs can exhibit distinct complexities (**Figure 1A-C)**, we delved further into the impact of different ELM classes (representing diverse motifs) on the performance of the MDTs and ATs. Initially, we examined the influence of the motif position within the peptides. To this end, we categorised the motifs based on their ELM classes’ regular expressions (**Supplementary Table S3**): N-terminal (N=4), C-terminal (N=15), or internal (N=139). The results revealed, as expected, that N- and C-terminal motifs were more amenable to accurate alignment compared to internal motifs (**Figure 3A,B**). This was particularly evident for alignment tools, where they consistently surpassed the MDTs.

**Figure 3.**
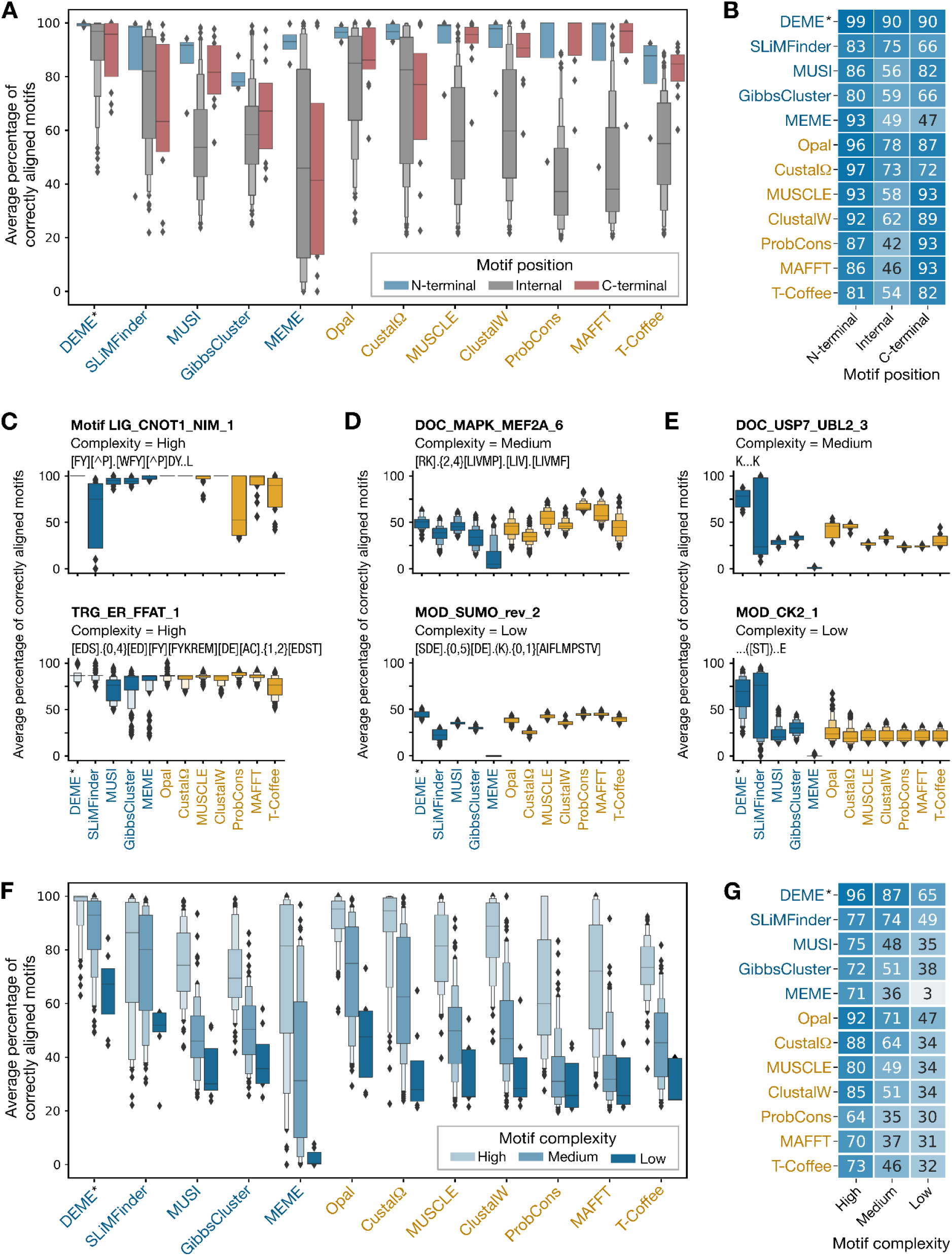
Effect of motif position and motif complexity on tools performances. **(A-B)** Effect of the motif position being N-/C-terminal or internal on the benchmarked tools performances by tool. The motif position refers to the ELM class definition of the motifs. Boxenplots **(A)** show the distribution of the percentages of correctly aligned motifs collapse at ELM class level while the heatmap **(B)** shows their average values. **(C-E)** Examples of motifs where all tools performed well **(C)**, poorly **(D)** or where MDTs outperformed ATs **(E)**. Each boxenplot shows all results collapsed at PSS, peptide lengths and noise percentage level. Results for all ELM classes can be found in **Supplementary Figure S6. (F-G)** Effect of motif complexity on MDTs and ATs performances. Boxenplots **(F)** show the distribution of the percentages of correctly aligned motifs while the heatmap **(G)** shows their average values. The results are collapsed at ELM class level, binned into 3 motif complexity tiers: High (motif probabilities of 0 to 10^−5^), medium (10^−5^ to 10^−3^) and low (10^−3^ to 1). *DEME results are the best results out of 11 runs using different expected motif length input values.

Among the motif diversity present in the 158 ELM classes explored in the current work (**Supplementary Figure S6**), we identified instances where all tools exhibited good performance in aligning internal motifs (**Figure 3C**). These cases included ELM classes DEG_CRL4_CDT2_1, DOC_MAPK_RevD_3, LIG_CNOT1_NIM_1, MOD_TYR_DYR, and TRG_ER_FFAT_1. These ELM classes share the characteristic of possessing multiple defined positions alongside minimal to no length variability. Conversely, certain ELM classes posed significant challenges for all tools (**Figure 3D**), such as DOC_MAPK_MEF2A_6, LIG_LIR_Gen_1, or MOD_SUMO_rev_2. These motifs exhibit a reduced number of defined positions and, in some cases, variable length gaps. Intriguingly, a subset of these challenging motifs demonstrated superior alignment performance using MDTs DEME and SLiMFinder compared to all other tools (**Figure 3E**). These ELM classes featured exceptionally short motifs, typically comprising only two defined positions and having fixed lengths, as exemplified by ELM classes DOC_USP7_UBL2_3, DOC_WW_Pin1_4, LIG_SH2_STAT3,

To conduct a more in-depth analysis of how motif complexity affects the performance of the benchmarked tools, we analysed the ELM classes probabilities as a descriptor of motif complexity. The ELM database provides motif probabilities for all ELM classes, defining the likelihood of a motif occurring by chance at a given position in the proteome (31). Generally speaking, the higher the probability of encountering a motif within a proteome, the lower its complexity and the more challenging it is to identify. As anticipated, we observed that lower complexity motifs were more difficult to align accurately by all tools (**Figure 3F,G**). Interestingly, when focused on the low complexity bin, only DEME achieved a mean surpassing 50% for the low probability motifs bin (65%), followed by SLiMFinder (49%) and then Opal (47%). Overall, all tools performed well on high complexity (or low probability) motifs. However, SLiMFinder exhibited instances where lower probability motifs were aligned suboptimally (**Supplementary Figure S7**).

Overall, all tools successfully identified the motifs for more straightforward cases, specifically N-/C-terminal motifs and high complexity motifs, but only three tools (DEME, SLiMFinder and Opal) produced good results across all motif classes tested.

### Discussion and conclusions

In this analysis, we established MEP-Bench (**Supplementary Data SD1**), a series of relevant datasets compiled from annotated instances in the ELM database to evaluate the performance of freely available motif discovery and sequence alignment tools. MEP-Bench datasets differ from previous approaches where artificially generated motifs were introduced into either randomly generated sequences following a particular distribution of probabilities (18) or into selected proteome regions (19). A total of 653,846 individual datasets were generated from 158 ELM classes, encompassing a range of peptide lengths, motif-containing peptide sizes, noise levels, and motif types with varying complexities. MEP-Bench sets represent a valuable resource for the motif community to benchmark future tools.

Our results indicated that increasing noise levels consistently reduced the accuracy of motif identification across all tools, emphasising the importance of noise reduction at the experimental or analytical stage prior to motif discovery. This approach is directly linked to the experimental system employed. Well-established methods to minimise non-specific binders include identifying peptides that appear across multiple independent experimental sets (referred to as “crapome” or “frequent flyers” typically applied to entire proteins), incorporating specific negative controls, or scoring peptides based on their experimental abundance (spectral counts in mass spectrometry, sequencing counts in next-generation sequencing) and/or their repetitive detection in replicated experiments. Additionally, increasing peptide length also diminished motif identification for all tools, albeit this parameter should be aligned with the research question and experimental limitations, considering that motifs can extend beyond 15 amino acids (e.g., ELM classes LIG_Actin_WH2_1 comprise 19 amino acids, and TRG_NLS_Bipartite_1 includes instances spanning 23 amino acids). With that said, it is still important to keep in mind that it is easier to identify motifs in shorter peptides, at least for the tools tested.

This raises the question of which tool provides the best performance. Our findings revealed that among the 12 benchmarked tools (5 motif discovery tools and 7 alignment tools), three emerged as the most effective: The DEME and SLiMFinder motif discovery tools, and the Opal alignment tool, though all of them had their limitations when dealing with low complexity motifs. DEME’s input requirements are also highly specific, requiring a negative control and the expected motif length. The output generated by the tools also plays a significant role, as alignment tools may correctly align motifs but fail to identify them for the user. In this regard, MDTs generating logos or regular expressions hold greater utility. Moreover, motifs can exhibit variable lengths, and not all tools are equipped to handle this variability. MEME, for instance, reports gapped motifs as two separate motifs, leaving it to the user to recognise and assemble the complete motif. In contrast, SLiMFinder’s algorithm can accommodate gapped motifs and GibbClusters even allows insertions or deletions in the input sequences (though this functionality was not evaluated in this study).

Another crucial consideration is the accessibility of a tool. While all tools provide access to their code and comprehensive documentation, this might not suffice for research groups lacking computational support. In such scenarios, a web-based tool can prove invaluable, neither DEME nor Opal offer one, and SLiMFinder’s web server is not particularly user-friendly. Among the evaluated MDTs, only MEME and GibbClusters offer user-friendly interfaces that grant access to the tools’ diverse options. SLiMFinder can also be accessed through PepTools (32), a peptide annotation web-tool that also provides motif discovery and analysis tools (8), albeit with the limitation that the input peptide sequences must belong to one of the species’ proteomes available in the tool. Additionally, both MEME and SLiMFinder are part of software suites specifically designed for motif identification and analysis, though only the MEME Suite (33) boasts a comprehensive user interface for accessing all functionalities (34).

In conclusion this study finds DEME, SLiMFinder and Opal to be the best performing tools for finding motifs in short peptides generated via high-throughput experimental settings, though no tool was capable of finding all motif types. If anything, these results show that there is still room for improvement when it comes to motif discovery either from a tool performance point of view or a needed accessible interface to make the tools more readily available for researchers.

## Supporting information

Supplementary Figures and Supplementary Tables overview

## Acknowledgements

We want to thank Dr. Abdellali Kelil and Dr. Stephen W Michnick for the help provided regarding MotifHound. We thank Hazem M. Kotb and Dr. Izabella Krystkowiak for critically reading the final manuscript.

## Availability of data and materials

The **Supplementary Table S2** compiling all results and the MEP-Bench datasets (**Supplementary Data SD1)** are available for download in the Zenodo repository (https://doi.org/10.5281/zenodo.10467208).

## List of abbreviations

AT: Alignment Tool
ELM: Eukaryotic Linear Motif (database)
MDT: Motif Discovery Tool
MEP-Bench: Motif Extraction from Peptides Benchmarking (datasets)
PSS: Positive Set Size
PSSM: Position-Specific Scoring Matrix
SLiM: Short Linear Motif

