## Supplementary Figures and Supplementary Tables overview for "Benchmarking computational tools for de novo motif discovery"

**A**

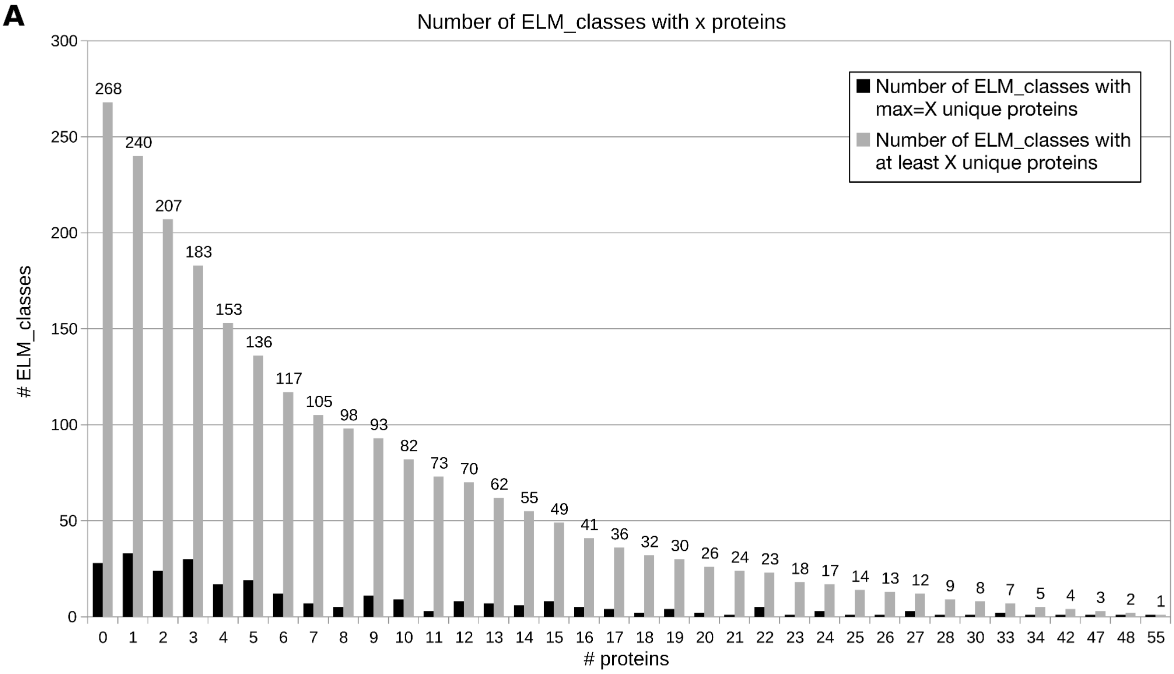


**B**

| Positive Set Size (PSS) | Peptide length in amino acids | | | | | | Noise percentage | | | | | | | |
|---|---|---|---|---|---|---|---|---|---|---|---|---|---|---|
| | 12 | 16 | 20 | 24 | 28 | 32 | 0 | 10 | 20 | 50 | 100 | 200 | 400 | 800 |
| 5 | | 156 | | | | | OK | SKIP | OK | 40% | OK | OK | OK | OK |
| 10 | | 94 | | | | | OK | OK | 25% | OK | OK | OK | OK | OK |
| 15 | 59 | 60 | 63 | 63 | 62 | 60 | OK | OK | OK | OK | OK | OK | OK | OK |
| 20 | 34 | 35 | 36 | 35 | 35 | 35 | OK | OK | OK | OK | OK | OK | OK | OK |
| 30 | | 8 | | | | | OK | OK | 23% | OK | OK | OK | OK | OK |
| 40 | | 4 | | | | | OK | OK | 25% | OK | OK | OK | OK | SKIP |
| 50 | | 1 | | | | | OK | OK | 24% | OK | OK | OK | OK | SKIP |

**Supplementary Figure S1 - ELM diversity influenced the generated MEP-Bench datasets.** Not all ELM classes contain the same number of instances, which means that not all sets of parameters explored in the current work produced the same amount of datasets. Because we enforced that only one instance could come from the same protein (to avoid generating overlapping peptides), the number of individual proteins in each ELM class defined the maximum theoretical number of datasets that could be generated for each PSS value. **(A)** The distribution of the number of ELM classes vs the number of proteins their instances come from is shown (as of February 2020). Black bars show the number of ELM classes that contain exactly that number of proteins while the grey bars show the cumulative number of ELM classes that have at least the given number of proteins. **(B)** The number of ELM classes available for dataset generation are shown (left, green section) relative to the values of the evaluated parameters (PSS and peptide length) used in the current work. The noise percentages used (right, grey section) depend on the PSS values and the total number of available instances of other ELM classes. Percentages that were skipped (red) or tweaked (yellow) are shown.

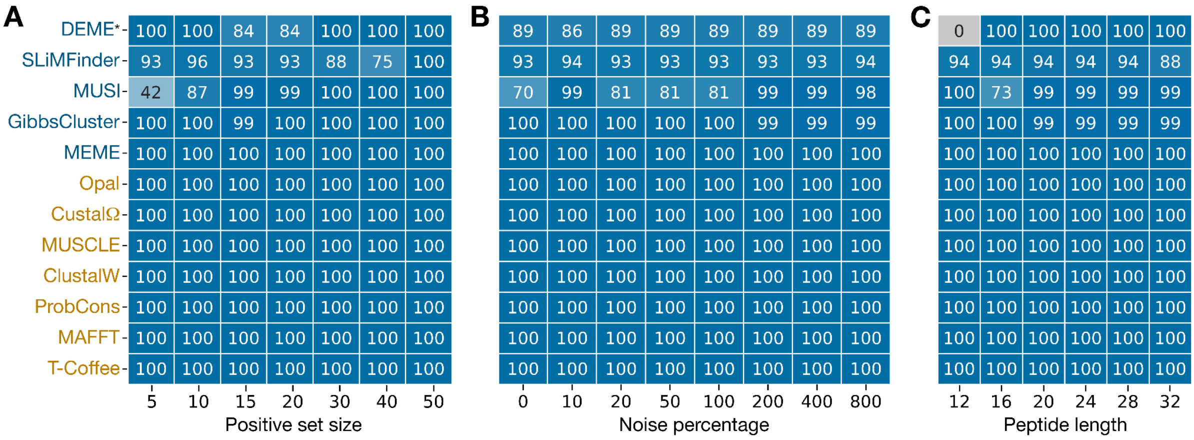


**Supplementary Figure S2 - Successful runs percentages.** Effect of the PSS **(A)**, noise percentage **(B)** and peptide length **(C)** on the percentage of successful runs per tool on all built datasets. *DEME results are for 11 runs using different expected motif length input values. The noise percentage groups for 20% and 50% also contain the tweaked values shown in **Supplementary Figure 1B**.

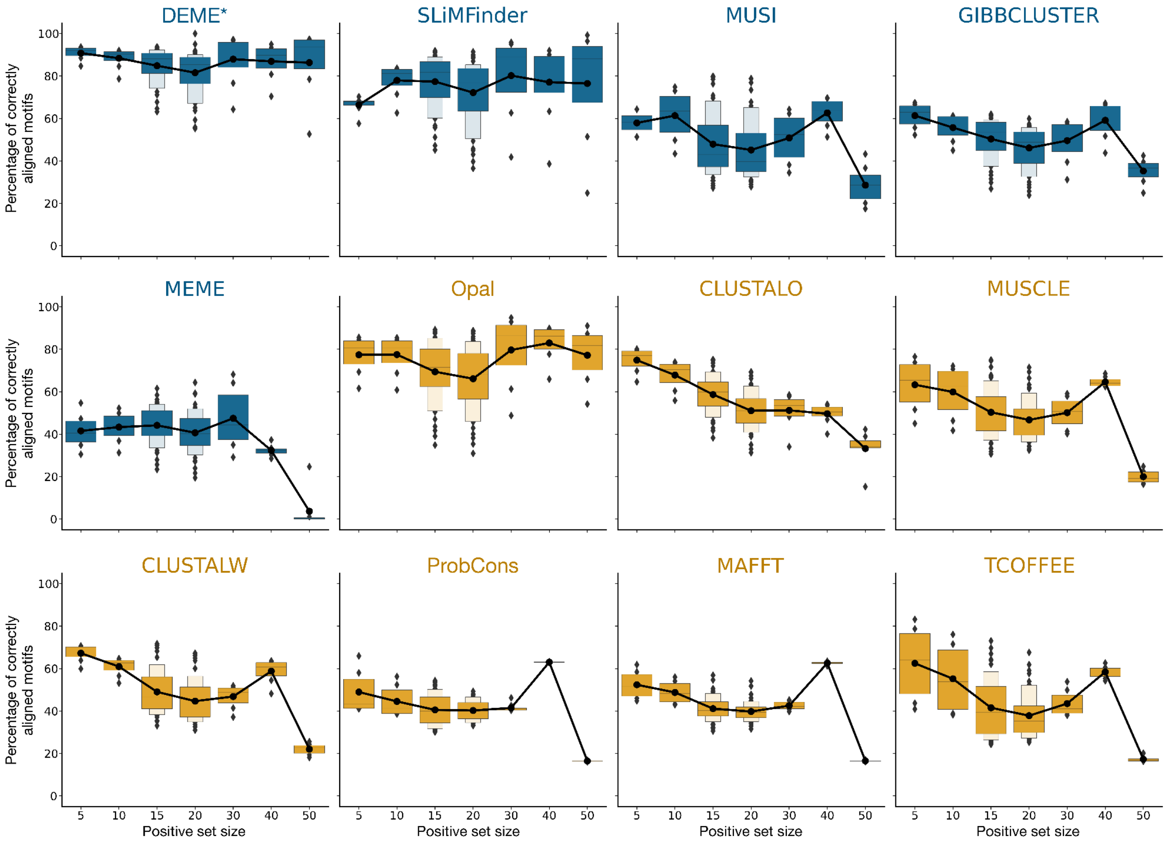


**Supplementary Figure S3 - Effect of the positive set size (PSS) on benchmarked tools.** Overall results showing the effect of the PSS on the percentage of correctly aligned motif results by tool. Boxenplots show the collapsed results at noise percentage and peptide length level while the overlaid black dot and line plot shows the mean value calculated over all non-collapsed data. *DEME results are the best results out of 11 runs using different expected motif length input values.

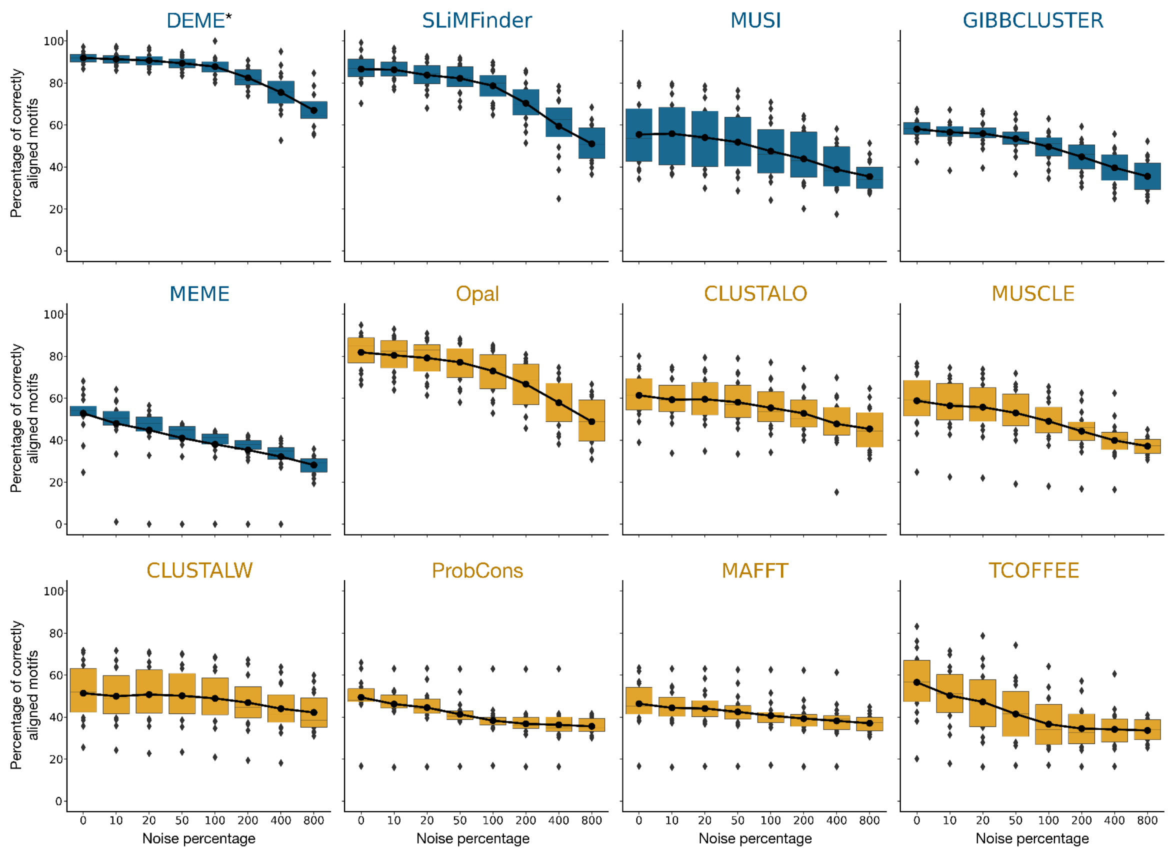


**Supplementary Figure S4 - Effect of the noise percentage on benchmarked tools.** Overall results showing the effect of added noise (as a percentage of the PSS value) on the percentage of correctly aligned motif results by tool. Boxenplots show the collapsed results at PSS and peptide length level while the overlaid black dot and line plot shows the mean value calculated over all non-collapsed data. The noise percentage groups for 20% and 50% also contain the tweaked values shown in **Supplementary Figure 1B**. *DEME results are the best results out of 11 runs using different expected motif length input values.

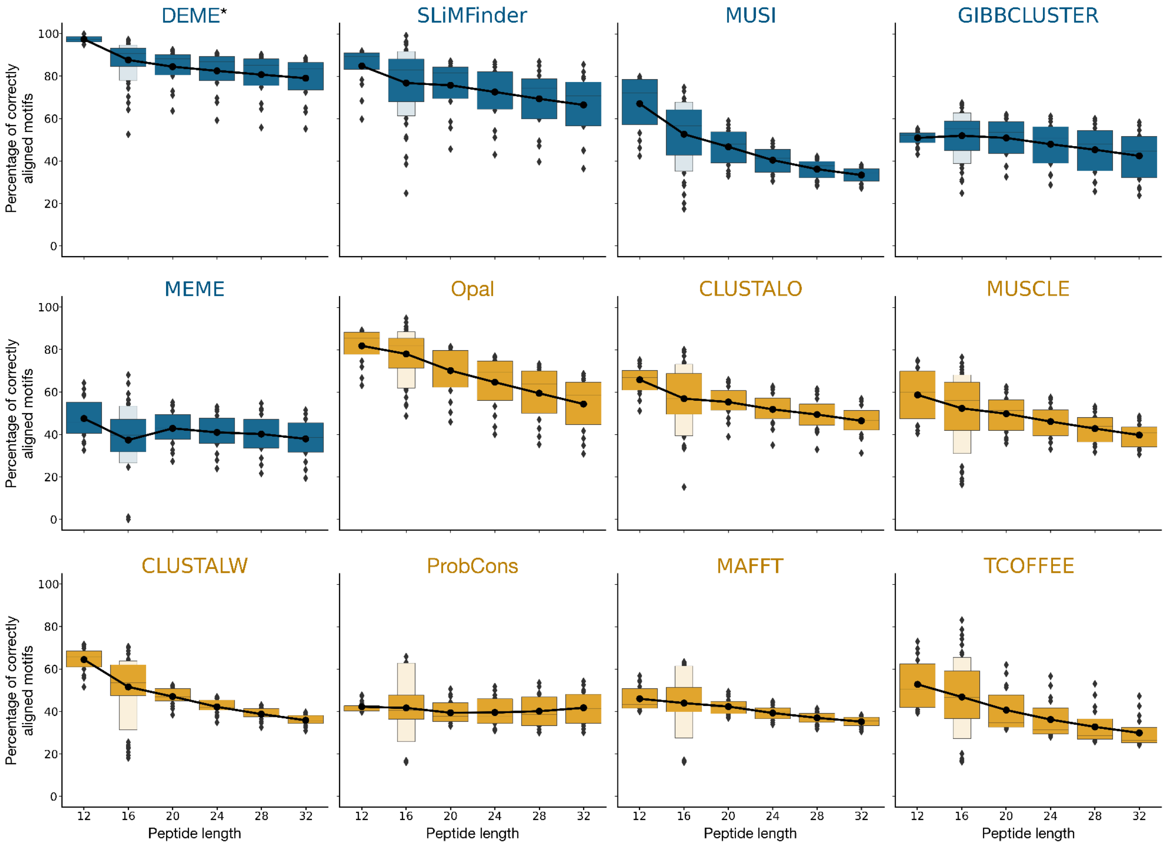


**Supplementary Figure S5 - Effect of the peptide length on benchmarked tools.** Overall results showing the effect of the generated peptides lengths (in amino acids) on the percentage of correctly aligned motif results by tool. Boxenplots show the collapsed results at PSS and noise percentage level while the overlaid black dot and line plot shows the mean value calculated over all non-collapsed data. *DEME results are the best results out of 11 runs using different expected motif length input values.

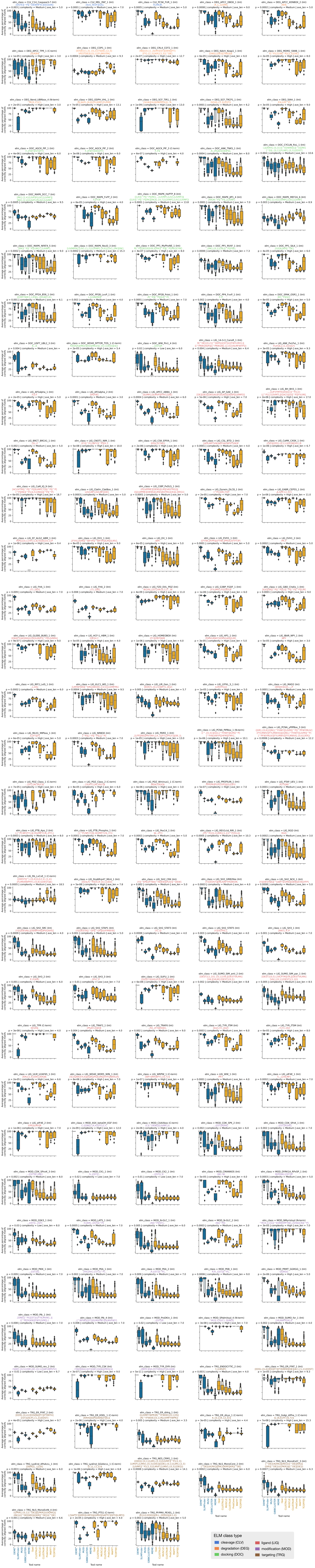
ELM class type
cleavage (CLV)
ligand (LIG)
degradation (DEG)
modification (MOD)
docking (DOC)
targeting (TRG)
Tool name

**Supplementary Figure S6 - General results by ELM class.** Overall results of the performance of all benchmarked motif discovery tools (blue) and alignment tools (orange) by ELM class. For each ELM class its position (N-/C-terminal or internal), regular expression definition, motif probability, complexity bin and average motif instance length are shown. Each boxenplot shows all results collapsed at PSS, peptide lengths and noise percentage level. *DEME results are the best results out of 11 runs using different expected motif length input values.

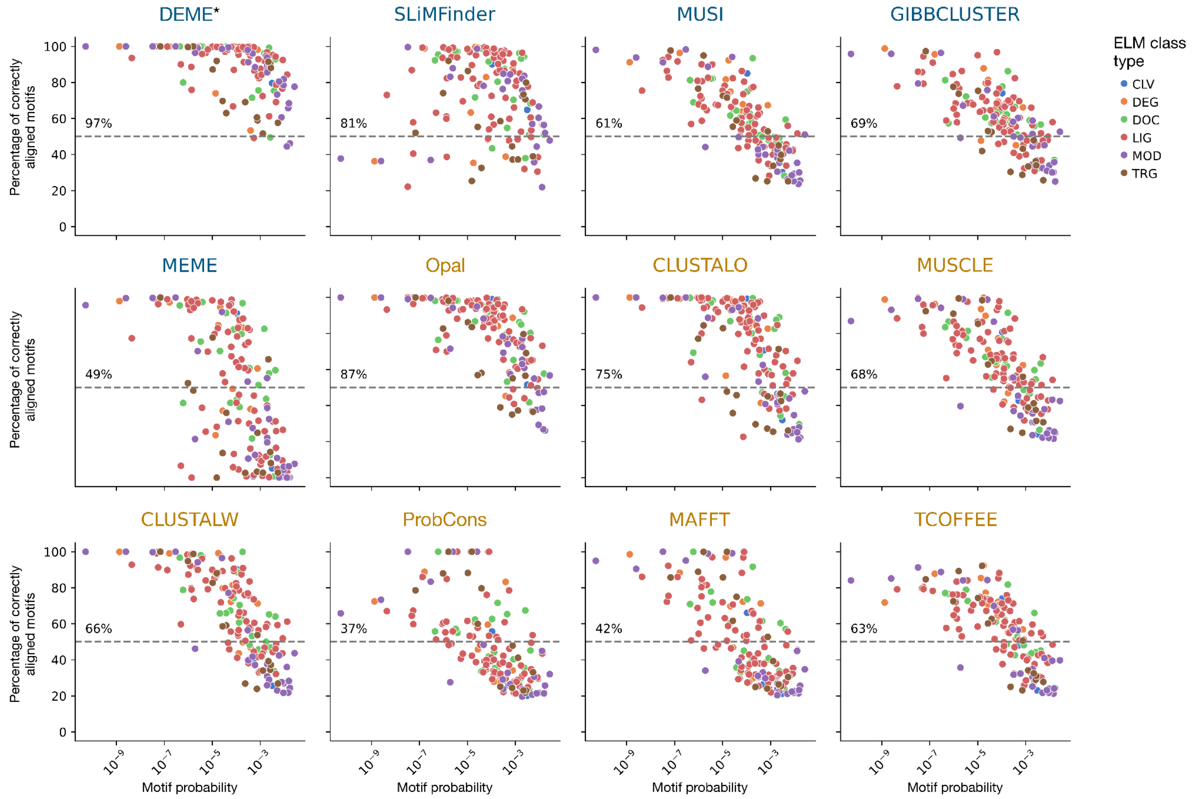


**Supplementary Figure S7 - Effect of motif probability on benchmarked tools.** Scatterplot showing the effect of motif probability on the average performance of the benchmarked tool. Dots are coloured based on the ELM class group. The percentage shown on the left of each plot represents the percentage of ELM classes for which the respective tool achieved an average of correctly aligned motifs above 50% (dotted line). Data is collapsed at ELM class level.

### OVERVIEW OF SUPPLEMENTARY TABLES AND DATA

**Supplementary Table S1 - ELM classes overview.** ELM Classes (as of February 2020) used for the generation of MEP-Bench datasets. The number of instances per class and proteins they come from is shown, together with their motif Regular Expression (RegEx) and probability. The position of the motif in the protein is classified as C-terminal (C), internal (I) or N-terminal (N) based on their RegEx.

**Supplementary Table S2 - Benchmarking results.** Compiles all tools run results for all MEP-Bench dataset replicates. For each replicate its ID (form 00 to 99), ELM class, PSS, noise peptides number and percentage and peptide length are shown. For the tools run on each dataset the run time and percentage of correctly aligned motifs are shown together with a regular expression or consensus of the motif found if the tool generated it. Due to its size, this table is available for download in the Zenodo repository (https://doi.org/10.5281/zenodo.10467208).

**Supplementary Table S3 - Benchmarking results, motif level analysis.** Motif (ELM class) level analysis results by tool. These results are collapsed at ELM classes level by averaging the percentage of correctly aligned motifs for all replicates, positive set size, noise percentage and amino acid length datasets by tool. The number of datasets contributing to the average is shown (N), together with the ELM classes regular expression, motif ELM instances average length and position, and the motif probability and complexity tier.

**Supplementary Data SD1 - Motif Extraction from Peptides Benchmarking (MEP-Bench) Sets.** All generated dataset files are available for download in the Zenodo repository (https://doi.org/10.5281/zenodo.10467208).
